# A generalised method for experiment design and model selection in the Bayesian framework

**DOI:** 10.1101/2023.10.24.563782

**Authors:** Prem Jagadeesan, Karthik Raman, Arun K Tangirala

## Abstract

Computational modelling of dynamical systems often involves many free parameters estimated from experimental data. The information gained from an experiment plays a crucial role in the goodness of predictions and parameter estimates. Optimal Experiment Design (OED) is being used to choose an experiment containing maximum information from a set of possible experiments. This work presents a novel Bayesian Optimal Experiment Design principle for generalised parameter distributions. The generalization is archived by extending the *β*-information gain to the discrete distributions. The *β*-information gain is based on what is known as the Bhattacharyya coefficient. We show that maximising the *β*-information gain is equivalent to maximising the angle between the prior and posterior distributions, reducing the posterior’s uncertainty. Further, we apply the proposed BOED criteria for two realistic experiment designs in systems biology. Firstly, we use the *β* information gain to choose the best measurement method for parameter estimation in a Hes1 transcription model. The measurement method selected by the *β*-information gain results in the minimum mean square error of the parameter estimates. In the second case, we employ the proposed information gained to select an optimal sampling schedule for HIV 1 2 LTR model. The sampling schedule chosen by the presented method reduces both prediction and parameter uncertainty. Finally, we propose a novel method for model selection using *β* information gain and demonstrate the working of the proposed method in the model selection in compartmental models.

## 1 Introduction

System identification of biological processes poses numerous challenges at each stage of system identification [1]. Experiment design, model structure selection and parameter estimation are three crucial stages in system identification [2]. The end goal of any modelling exercise is a *useful* model; in modelling biological processes, the usefulness is generally assessed by the quality of predictions, parameter estimates and interpretability. The quality of the parameter estimates and predictions largely depends on the data set’s information content and the model structure’s nature. Obtaining precise/practically identifiable parameter estimates from noisy and limited data is one of the prevailing challenges in systems biology. Further, the nature of the multi-parameter non-linear models being sloppy hampers the quality of the parameter estimates [3, 4, 5, 6]. Anisotopic sensitivity in the parameter space is the prime reason. It is well known that the factors contributing to the sloppiness are both experimental conditions and the nature of the model structure [4, 7, 8]. Hence, maximising the information content in the experimental data is an ineludible solution to obtain good predictions and parameter estimates.

Fisher Information Matrix (FIM) and covariance matrix of the parameter estimates are the most commonly used Optimal Experiment Design (OED) criteria [9, 10]. Following are the three frequently used optimal design criteria to minimise variability in the parameter estimates (i) *A* − *optimal*, maximising trace the trace of the FIM, (ii) *E* − *optimal*, maximising the minimum eigenvalue (iii) *D* − *optimal*, maximising the determinant of the FIM. Some works based on optimising FIM-based criterion are [11, 12, 13]. This work focuses on maximising the information gained in the Bayesian framework. One of the perceived advantages of the Bayesian framework is its ability to incorporate the prior information about a parameter in the form of a p.d.f; moreover, posterior parameter distributions obtained from Bayesian inference allows us to use global methods rather than the aforementioned local Fisher information-based methods as the Fisher information for nonlinear predictors is usually numerically computed at the optimal parameter estimates.

Bayesian Optimal Experiment Design has seen several developments in the past two decades. A concise review of BOED is given in [14, 15]. Initial attempts were focused on reducing the information entropy in the posterior parameter distribution, as entropy is a measure of uncertainty [16, 17, 18]. Juliane *et al*. use the concept of entropy and mutual information to design experiments that maximise the information content in terms of both parameter estimates and predictions [19]. BOED based on reducing the prediction variability was proposed in [20]. A computationally efficient method is proposed in [21]. A decision-theoretic approach with Kullback-Leibler divergence as the design criterion was proposed to design experiments for non-linear systems [22]. A recent FIM-based method to choose experiments to optimise the confidence region of the parameter estimates has been proposed in [23].

The problem of model selection is also linked to the information contained in the data set. The problem of model selection and the problem of optimal experiment design are two sides of the same coin. In the problem of model selection, the experimental data is fixed, and the central idea is to choose the model that absorbs maximum information in the data. In contrast, in the optimal experiment design, it is *vice versa*. Akaike information criterion (AIC) and Bayesian information criterion (BIC) are the most commonly used model selection criterion. They attempt to strike a balance between the goodness of fit and parsimony of models [2]. However, they do not guarantee the goodness of the parameter estimates. To circumvent this issue, an experiment design solely for model selection was proposed in [24]; they used the experiment design criterion for Jensen–Shannon divergence. Daniel *et al*. proposed a Hellinger distance-based experiment design for model selection [25].

In all of the aforementioned information criteria for experiment design and model selection, one of the two following challenges prevails (i) Boundedness, (ii) Interpretability of bounds in terms of parameter estimates. From a system identification perspective, bounded information measures and the interpretability of the bounds have significant utility. Also, assuming joint prior and posterior parameter distributions to be joint Gaussian is a strong assumption in most cases. Hence, in this work, we extend the *β*-information gain-based BOED proposed in [26] to non-Gaussian prior and posterior distributions. In addition to the advantages proposed above, estimating the Bhattacharyya coefficient is computationally easier than the Kullback-Liebler divergence [27]. We work with the discretised sample prior and posterior distributions. The *β*-information gain proposed is bounded and has a natural interpretation in terms of the precision of the parameter estimates [26]. The *β* information gain is based on what is known as a Bhattacharyya coefficient, and Bhattacharyya distance [28]. The Bhattacharyya distance measures the distance between two probability distributions, while the Bhattacharyya coefficient can be interpreted as the amount of sample overlap between two distributions. Further, we show that maximising the *β* information gain is equivalent to maximising the angle between prior and posterior distributions resulting in a reduction in posterior uncertainty. We propose a machine learning-based method to estimate the *β*-information gain for discrete distributions.

We demonstrate the proposed BOED on two benchmark systems biology problems, (i) select a measurement system for optimal parameter estimation in Hes 1 model, the measurement method selected by the proposed method resulted in the minimum mean square error of the parameter estimates, (ii) select an optimal six-point sampling schedule for parameter estimates in HIV patients under treatment intensification, the parameters obtained from the sampling schedule resulted in minimum prediction and parameter uncertainty. Finally, we propose a novel method for model selection based on the *β*-information gain and demonstrate the working of the method in two-compartmental pharmacokinetic models.

The rest of the paper is organised as follows: Section 3 contains the preliminary concepts and definitions. Section 4 presents the preliminary results on extending the *β*-information gain to discrete distributions and its application to experiment design and model selection and illustrates the working of the proposed method with numerical examples. Section 5 contains the method for estimating the *β*- information gain and model selection. The paper ends with concluding remarks in section 6.

## 2 Preliminaries

### 2.1 Experiment design problem in dynamical systems

Consider a general non-linear ODE model with discrete time noisy measurements. The mathematical description of the model is given in 1.

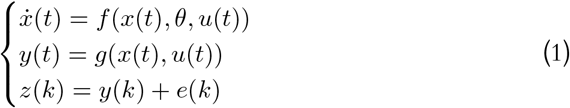

The structure of the state equation is assumed to be known. The unknown variables are the parameter vector *θ*, the input *u*(*t*), the noise characteristics of *e*(*k*), the number of data points *N*, and the sampling interval *t*_*k*_. In some cases, the structure of *g*(*t*) and the way the input enters the model are also unknown. The goal of the experiment design is to design one or more experiments to estimate the parameters *θ* from data based on optimal criteria.

#### 2.1.1 Degrees of freedom for Optimal Experiment Design of dynamical systems

The goal of any optimal experiment design problem is to select the experiment that has maximum information content. Information supplied by an experiment is a function of the quality and quantity of data [18]. In dynamical system formulation, factors affecting the quantity and quality of data are given in Fig. 1.

**Figure 1:**
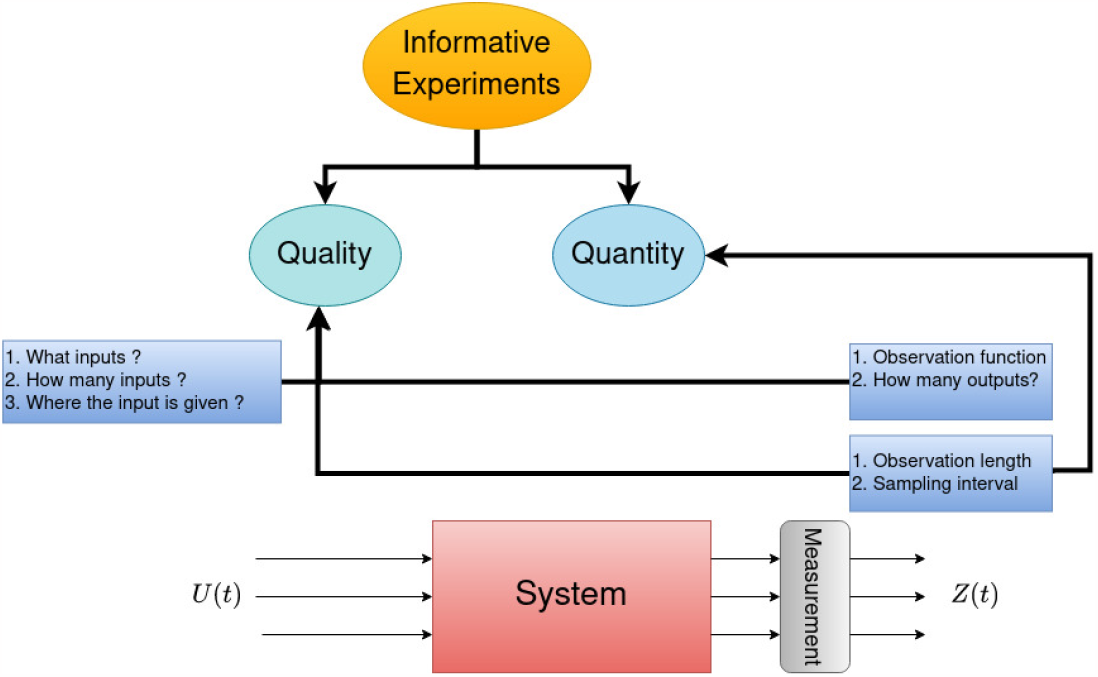
Degrees of freedom in the experiment design of dynamical systems

**Figure 2:**
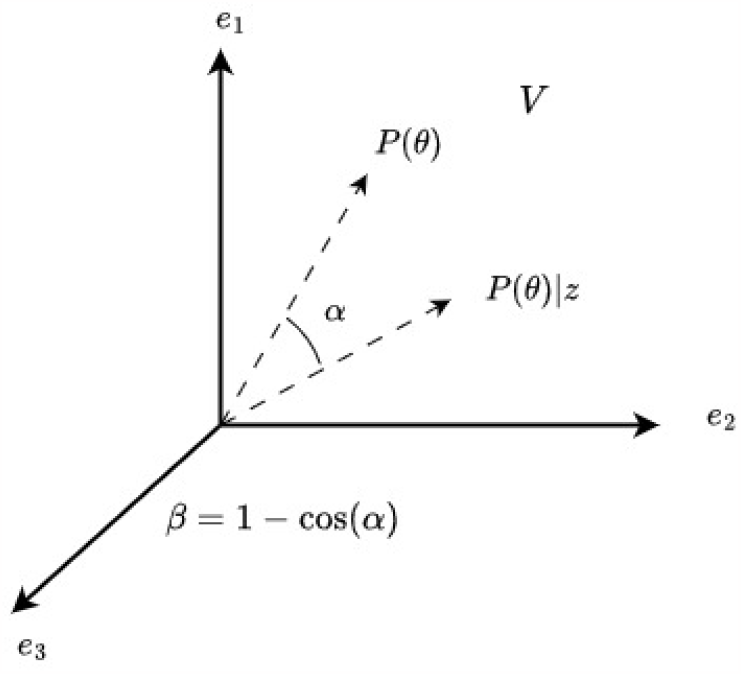
Three-dimensional view of the vector space *V*

The quality of data is affected by both input and output variables; from input direction, the number of inputs, the nature of input functions and the way the input enters the system affect the data quality. From the output direction, the number of outputs measured, signal-to-noise ratio, nature of the output function, number of data points, duration of the output and sampling interval affect the data quality. The quantity of data is affected by the number of data points, sampling interval and duration of the measurement.

### 2.2 Bayesian optimal experiment design

In the Bayesian framework, OED is defined as follows: given a set of discrete experimental conditions *χ* = {*z*_1_ … *z*_*n*_}, prior information of the parameters in the form of a probability distribution function and a model structure ℳ, choose an experiment *z*_*i*_ that maximises the distance between the prior and posterior of the parameters (2).

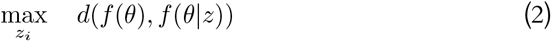

where *d*(·) any distance measure between probability distributions, *f* (*θ*) is the joint prior distribution of parameters, *f* (*θ*|*z*) is the joint posterior distribution of parameters. This class of Bayesian optimal design problem’s primary objective is to reduce the posterior distribution’s uncertainty. However, unlike FIM based metric, where the focus is only on reducing the variability in the posterior distribution, obtaining a posterior distribution much different (in terms of assumed statistical/probabilistic distance) from the prior distribution is informative not only from the perspective of reduction of variability but also from moving the MAP estimate close to the true parameters, if the data is informative.

### 2.3 Bhattacharyya coefficient and Bhattacharyya distance

Bhattacharyya distance (*B*_*d*_) is a measure of similarity between two statistical distributions [28]. The *B*_*c*_ can be used to quantify the relative closeness of two samples. The *B*_*c*_ of two densities *f*_1_(*θ*) and *f*_2_(*θ*) is given as

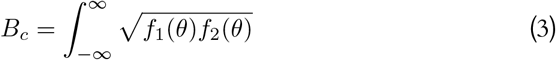

where *f*_1_(*θ*) and *f*_2_(*θ*) have identical outcome space. *B*_*c*_ is bounded between 0 ≤ *B*_*c*_ ≤ 1. A distance measure associated with this is the *B*_*d*_:

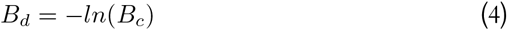

The Bhattacharyya distance is bounded between 0 ≤ *B*_*d*_ ≤ ∞. It is noteworthy that the *B*_*d*_ is not metric as it does not obey the triangle inequality.

In this work, we work with arbitrary discrete prior/posterior distributions where the closed expression for the distribution is unavailable; hence we turn towards the *B*_*c*_ for discrete distributions.

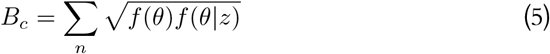

where *n* is the number of bins in the multivariate histogram. The goodness of the estimate of the discrete Bhattacharyya coefficient depends on the number of bins. The Bhattacharyya coefficient is widely used in feature extraction [29] and optimal signal selection [27].

## 3 *β*-information gain for generalized distributions

In our previous work [30], we showed that the *β*-information gain index could be used as a Bayesian Optimal Experiment Design criterion. We demonstrated that *β*-information gain not only reduces the uncertainty in the posterior but also prediction uncertainty. The primary assumption of the previous work is that the joint prior and posterior distributions are Gaussian. In this work, we extend the *β*-information gain to non-Gaussian prior and posterior distributions.

In the case of discrete distributions, the information gain is defined as

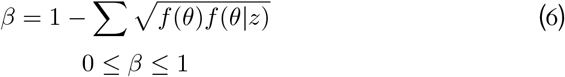

*Case 1:* If *f* (*θ*) = *f* (*θ*|*z*) then *β* = 0. No new information is present in the experimental data apart from the prior.

*Case 2:* In the case of discrete probability mass functions, if only one of the samples is accepted in the posterior and if that sample has negligible probability in the prior, then *β* ≈ 1. This indicates that the experiment is highly informative such that it has picked a low probability sample from the prior.

### 3.1 Geometric interpretation of the new information gain

Consider an *n*-dimensional vector space *V* where *n* is the cardinality of the outcome space of possible parameter values. Each element *v* of the vector space is the square root of probabilities. Hence, each vector can be considered as a transformed probability mass function.

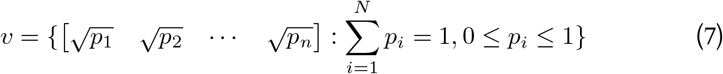

The domain of the vector space is restricted to,

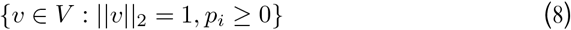

The basis of the vector space are orthonormal basis

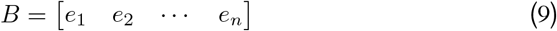

The basis vectors are p.m.f. of type Kronecker-delta function, i.e., the pmf has the form of Kronecker-delta function (Deterministic). Given a vector *v*, the angle that it makes with a basis vector is inversely proportional to the probability of the corresponding outcome.

In a Bayesian framework, the sampled prior and posterior probability density function can be represented as vectors in *V*. The proposed information gain index is

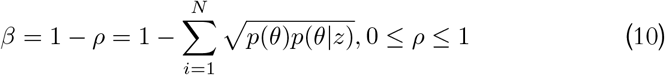

where *ρ* is the Bhattacharyya coefficient, the Bhattacharya coefficient can be seen as *cos* of the angle between two vectors in 𝒱. We also know that the Bhattacharyya coefficient quantifies the amount of overlap between the distributions; hence, the angle represents the amount of overlap. The lesser the angle between two vectors, the more they are similar.

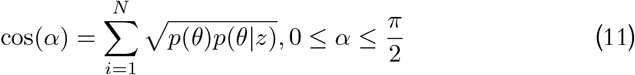

*Case 1:* When *α* = 0 ⇒ *ρ* = 1 ⇒ *β* = 0. There is no new information in the data. Prior and posterior are identical.

*Case 2:* When 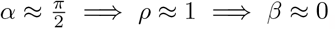. Data has improved prior to the maximum in terms of reducing uncertainty.

**Theorem 1**. *If the prior distribution of a parameter θ is uniform with countably infinite outcome space and posterior distribution in the form of Kronecker-delta for a given experiment e*_*k*_, *then in the space V*, *the angle between prior and the posterior distributions is* 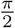 *resulting in maximum β information gain*.

*Proof*. The angle between the prior and posterior distribution (Bhattacharyya coefficient *B*_*c*_) is derived as follows,

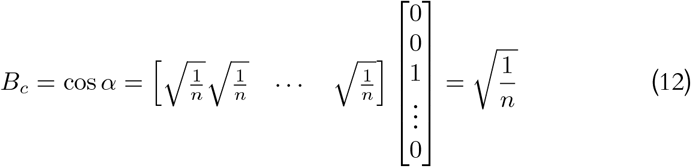

For arbitrarily large *δ*,

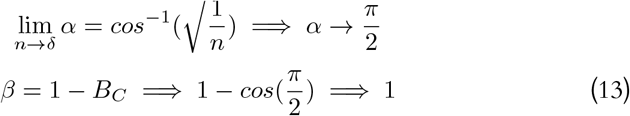

This concludes the proof □

Thus maximising the angle is equivalent to reducing the uncertainty in the prior. In all other cases, maximising the angle will drive the posterior vector towards one of the basis. The angle cannot be 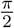 because prior either has to be in an infinite dimensional space or one of the basis as prior, which is not possible in practical scenarios.

### 3.2 Application in Model Selection

In the dynamical modelling of biological processes, model selection is considered one of the three classical problems from a system identification point of view [1]. Given a set of possible model structures, prior information on the parameters and a data set, the problem is to find the model that best explains the given data set. Factors influencing model selection are (i) the parameter estimates and (ii) the goodness of the predictions. The ensemble and asymptotic properties of the estimates generally assess the goodness of the estimates. However, minimising the uncertainty in the parameter estimates and prediction for small sample scenarios is considered sufficient. This work uses a new information gain-based approach for model selection in the Bayesian framework.

#### Problem Statement

Given a model set ℳ = {*M*_1_, *M*_2_,, *M*_*n*_}, a data set *Z*, and the priors of the parameters in the form of a p.d.f, choose an appropriate model structure that best explains the data.

#### Measure of Goodness of fit

Compute the sum-squared error for every *θ* in the posterior distribution

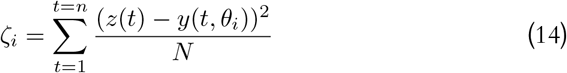

Then the measure of goodness of the predictions is computed by

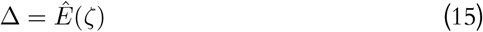

In this work, we use the sample mean as an estimator of *E*(*ζ*).

### 3.3 Numerical Results

#### 3.3.1 Hes 1 Oscillator

We consider a three-state model with mRNA regulatory dynamics of Hes 1 oscillator [31]. The state variable *m, p*_1_, and *p*_2_ represent the Hes1 mRNA concentration, cytoplasmic concentration and nuclear protein concentration. The parameter *P*_0_ is the amount of Hes1 protein in the nucleus, *v* is the translation of Hes1 mRNA, *h* is the hill coefficient, and *k*_1_ is the rate of transport of Hes1 protein. The model equations are given in (2)

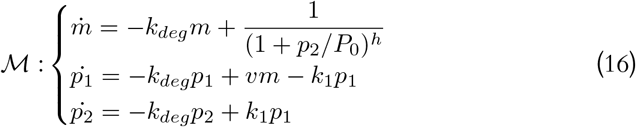

The parameter *k*_*deg*_ is experimentally measured as 0.03 min^−1^. In this case, using western bolts, it is possible to measure either the mRNA using real-time PCR or total Hes1 protein concentration (*p*_1_ + *p*_2_). In this example, we use the proposed method to investigate which mode of measurement results in minimum parameter uncertainty. The reference parameters of the model are *θ* = [1.4 5.7 0.002 0.009]. The initial conditions are *x*_0_ = [5 2.5 2.5] and the model is simulated for *t* = 0 to *t* = 200 seconds with sampling interval *T*_*s*_ = 5 seconds.

The parameters are estimated using Markov Chain Monte Carlo-based Approximate Bayesian Computation (MCMC-ABC). Fig. 3 shows the histogram of the prior and posterior distributions along with the actual parameters. The data has significantly shrunk the uncertainty in the parameters *P*_0_ and *h*. However, the parameter *v*_*m*_ has been estimated with poor precision. On the other hand, in the combined measurement of *p*_1_ and *p*_2_, the parameters *P*_0_ and *k*_1_ have been estimated with considerable precision, but the parameters *h* and *v*_*m*_ are estimated with poor precision. The information gain is maximum for the experiment where mRNA is measured. From Table 2, it can be observed that both bias and reduction in the variance of the parameter estimates. By measuring mRNA alone, three out of four parameterise are precisely measured while measuring two proteins. Only two parameters are precisely measured, and there is a significant bias in the parameter *h* and *P*_0_. In addition, from Table 3, the experiment with maximum information gain has minimum Mean Square Error (MSE).

**Table 1:**
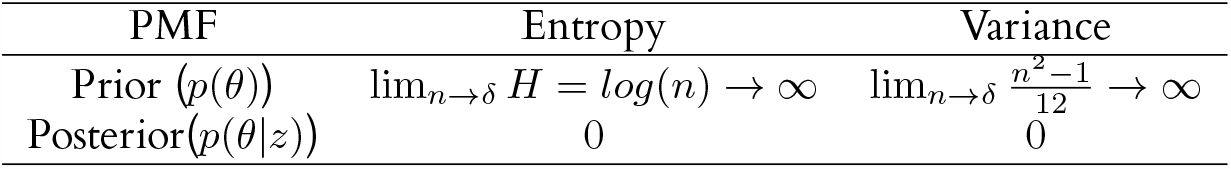
Entropy and Variance of the uniform prior and the Kronecker-delta posterior.

**Table 2:** Summary statistics of sample posterior distributions.

| Parameter | Estimate | | Bias | | Reduction in $\sigma_\theta^2$ | |
| --- | --- | --- | --- | --- | --- | --- |
| | $m$ | $p_1 + p_2$ | $m$ | $p_1 + p_2$ | $m$ | $p_1 + p_2$ |
| $P_0$ | 1.4066 | 1.3259 | 0.0066 | -0.0741 | 99.7% | 95.88% |
| $h$ | 5.6319 | 5.2985 | -0.0681 | -0.4015 | 96.5% | 32.47% |
| $v_m$ | 0.0211 | 0.0206 | 0.0011 | $5.7 \times 10^{-4}$ | 0.28% | 62.12% |
| $k_1$ | 0.0911 | 0.0897 | 0.0011 | $-3.1 \times 10^{-4}$ | 77.3% | 93.57% |

**Table 3:**
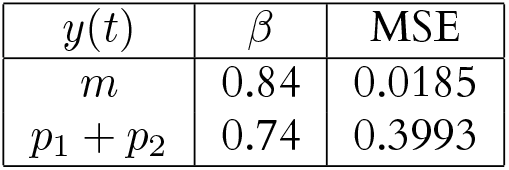
Information gain *β* and MSE for both the experiments.

| $y(t)$ | $\beta$ | MSE |
| --- | --- | --- |
| $m$ | 0.84 | 0.0185 |
| $p_1 + p_2$ | 0.74 | 0.3993 |

**Figure 3:**
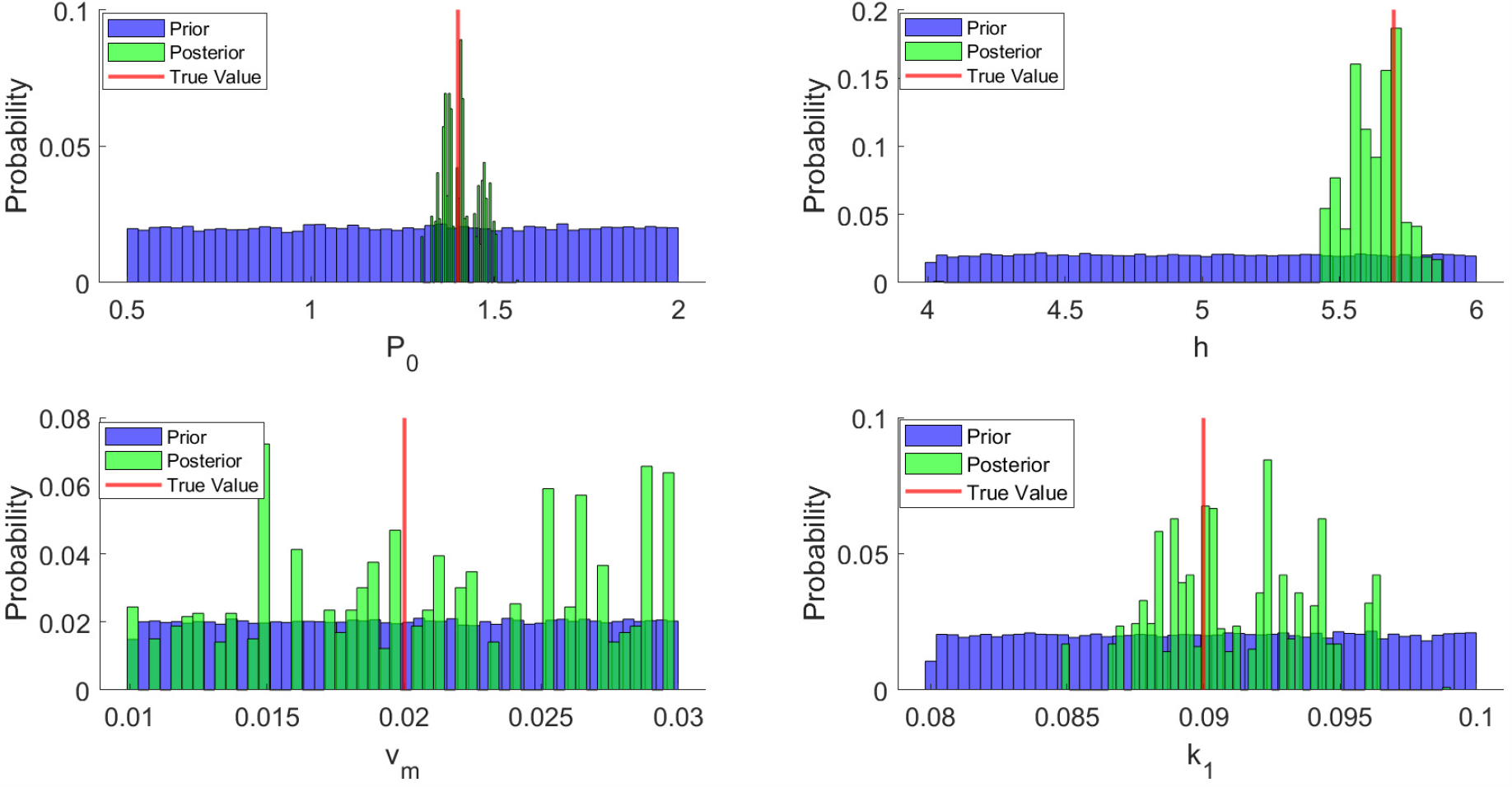
Prior and posterior distribution of all the parameters estimated by measuring mRNA. The variability in parameters *P*_0_ and *h* has considerably reduced compared to the parameters *v*_*m*_ and *k*_1_.

**Figure 4:**
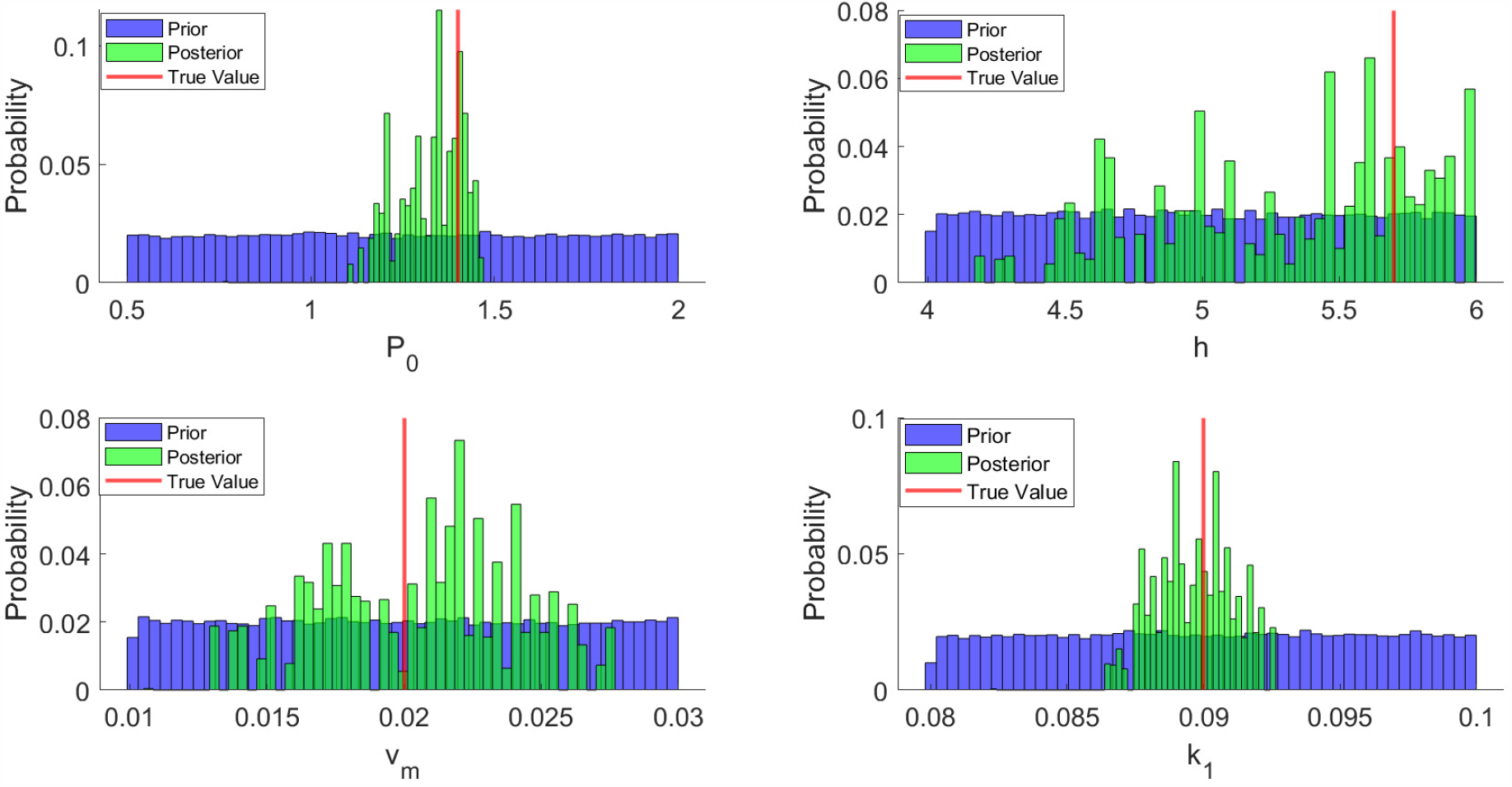
Prior and posterior distribution of all the parameters estimated by measuring both proteins. The variability in parameters *P*_0_ and *k*_1_ has considerably reduced compared to the parameters *v*_*m*_ and *h*.

This case study shows that the experiment chosen by *β* information gain has minimum mean square error. Further, it reduces both parameter uncertainty and bias for the stiff parameter estimates.

#### 3.3.2 HIV 1 2-LTR Dynamics

In this example, we design optimal experiments to select optimal sampling times for estimating model parameters in the HIV 2-LTR model. This model is developed in [32] to predict 2-LTR concentration post-treatment intensification with an integrase inhibitor. The clinical trials are primarily costly, and along with it, the burden experienced by the patients must also be considered. Institutional Review Board (IRD) regulates clinical trials on patients. Hence optimal scheduling becomes extremely important in clinical trials. The model equations are given below:

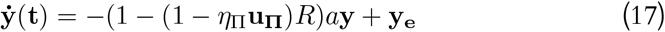

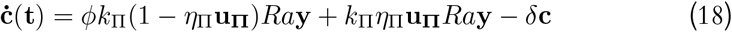

where **c** is a concentration of 2-LTR circles and **y** is a concentration of actively infected CD4+ T-cells, **y**_**e**_-exogenous sources of infected cells unaffected by treatment, *ϕk*_Π_(1 − *ν*_Π_**u**_**Π**_)*Ra***y** - rate at which 2-LTR circle forms post intensification, (1 − (1 − *ν*_Π_**u**_**Π**_)*R*)*a***y**- turnover rate of the infected cells completing a cycle, **u**_**Π**_ is a binary parameter indicating the presence or absence of the treatment and *δ* being the decay rate of the 2-LTR circles’ formation. From [32], it is assumed that the dynamics have attained steady-state prior to treatment intensification; the steady-state equations are

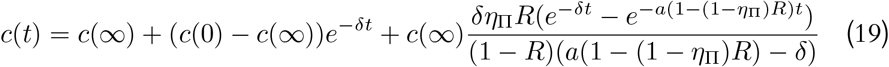

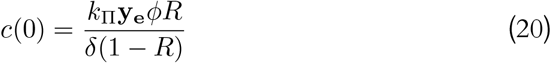

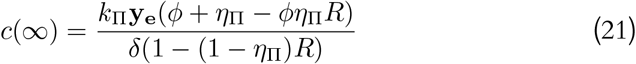

##### Experimental conditions

In this work, we replicate the experimental conditions used in [32]. We consider the production of 2-LTR circles in the presence of 2-LTR circle treatment intensification. The replication of the 2-LTR circles in the presence of treatment is quite high, as predicted by the model and has been clinically observed in doi:10.1098/rsif.2013.0186. The measurement technique used to measure 2-LTR circles was a PCR process which introduces log-normal uncertainty; the measurement equation, along with measurement noise, is given below.

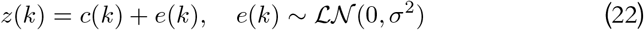

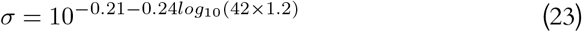

The variance of the noise in the (11) is taken from [32], which replicates the measurement error induced by a PCR process. The parameter values used for data generation are given in Table 2.

This work considers four different six-point sampling schedules obtained from four different optimal design criteria [33]. We investigate four sampling schedules informative with respect to *β* information index. We use MCMC-ABC to estimate parameters. We generate two hundred noise realisations with the same SNR and estimate *β*-information gain.

From Table 5, we can see that the average *β*-information gain is maximum for the *D*-optimal sampling schedule. Followed by *E*-optimal, *A*-optimal and *ELK*- optimal. From Fig. 5 and Table 5, it is evident that all the estimates of *β* are tightly constrained. The mean square error is also the minimum for the *D*-optimal sampling schedule.

**Table 4:**
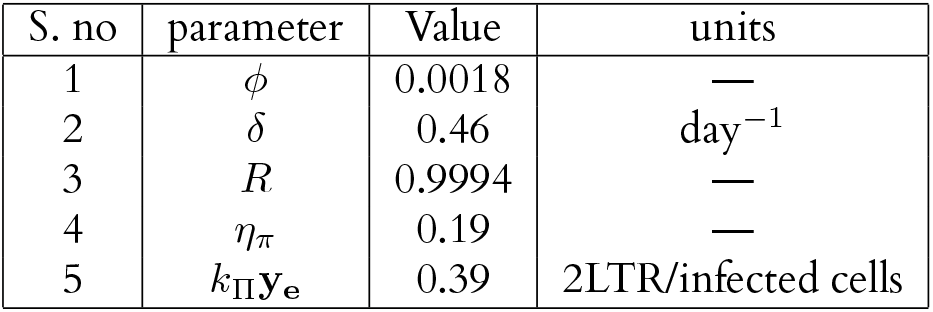
Parameter values estimated from the experiment.

**Table 5:** Sampling times selected from different optimal criteria.

| Experiment id | Optimal criterion | 1 | 2 | 3 | 4 | 5 | 6 | $\hat{E}(\beta)$ | MSE |
| --- | --- | --- | --- | --- | --- | --- | --- | --- | --- |
| Ex1 | $A$-optimal | 0 | 1 | 4 | 9 | 27 | 28 | 0.859($\pm 0.0006$) | 0.65 |
| Ex2 | $D$-optimal | 0 | 1 | 2 | 4 | 11 | 26 | 0.893($\pm 0.0008$) | 0.54 |
| Ex3 | $E$-optimal | 0 | 2 | 4 | 10 | 26 | 27 | 0.867($\pm 0.004$) | 0.69 |
| Ex4 | ELKD | 1 | 2 | 3 | 11 | 27 | 46 | 0.846($\pm 0.006$) | 0.745 |

**Figure 5:**
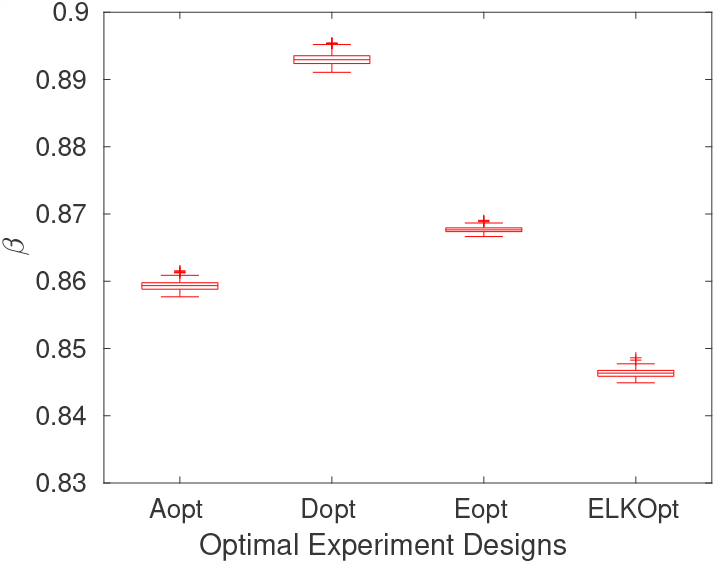
*β* information gain for each sampling schedule

From Table 6, all the sampling schedules are informative with respect to the parameter *R* (probability of infected cells infecting a target cell). *D*-optimal schedule is more informative with respect to the parameter *ϕ* (Ratio of probability of 2-LTR circle formation) and *δ* (decay rate of 2-LTR circles). With repect to the parameters *η*_*π*_ and *k*_*π*_*y*_*e*_, *E*-optimal sampling schedule contains more information.

**Table 6:** Reduction in variance for all the parameters.

| S. no | Parameter | A-opt | D-opt | E-opt | ELK-opt |
| --- | --- | --- | --- | --- | --- |
| 1 | $\phi$ | 38.7% | 58.0% | 39.5% | 25.9% |
| 2 | $\delta$ | 80.9% | 84.8% | 80.6% | 47.8% |
| 3 | $R$ | 99.7% | 98.2% | 99.9% | 99.1% |
| 4 | $\eta_\pi$ | 47.2% | 14.7% | 63.0% | 11.2% |
| 5 | $k_{\Pi}y_e$ | 9.5% | 22.3% | 69.3% | 29% |

From Fig. 6, it is clear that the samples obtained from *D*-optimal sampling schedule constraints the predictions most than all other sampling schedules. The *EKLD*-optimal sampling schedule having the least information gain has poor prediction uncertainty. The *β*-information gain has chosen the experiment (*D*-optimal) with the minimum mean square error. Further, the shared parameters *ϕ, α* and replication rate (*R*) (crucial for detecting an ongoing infection) has been estimated with very high precision in the chosen experiment.

**Figure 6:**
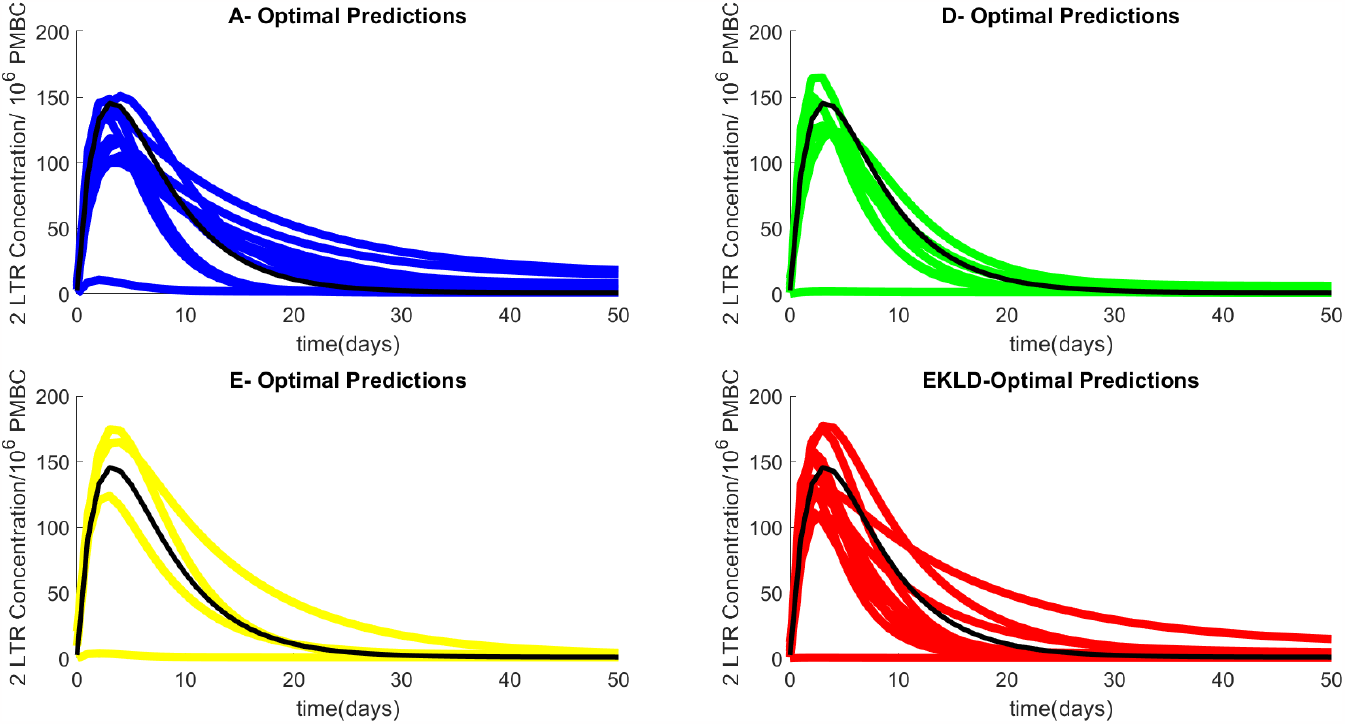
Predictions from the sample posterior parameter distributions for all the sampling schedules. Predictions from *D*-optimal are well constrained around the predictions from true parameters. Black-Predictions from true parameters, Blue-Predictions from A-optimal, Green-predictions from D-optimal, Yellow- predictions from E-optimal, Red-predictions from EKLD-optimal, Black-True Predictions

**Figure 7:**
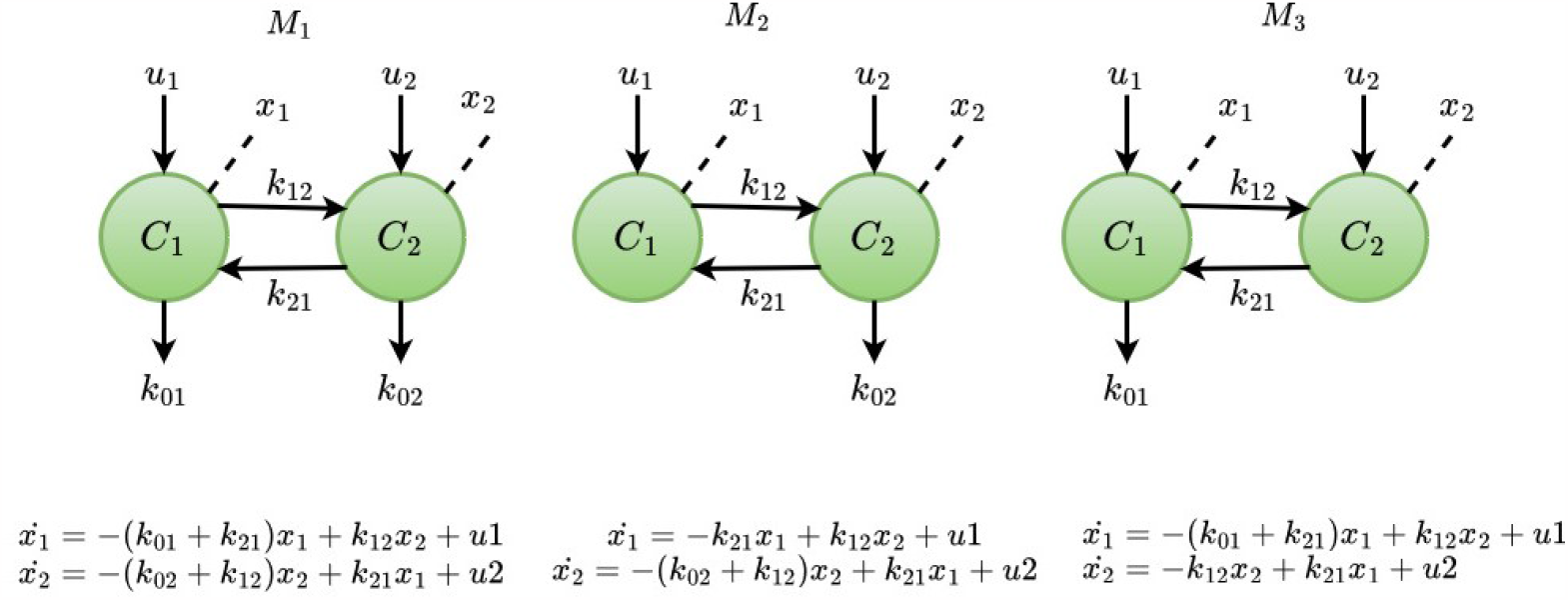
Candidate models

#### 3.3.3 An illustrative example on model selection

In this example, we demonstrate the working of the proposed method in the leak ambiguity problem in two compartmental multi-input, multi-output models given in [34]. The data generating process is (*M* 1) with nominal parameter vector *θ*^*^ = [1 1.5 2 1]^*T*^. The output is measured as *z*(*t*) = *x*_1_(*t*) + *x*_2_(*t*) + *e*(*t*). The noise variance is adjusted such that the signal-to-noise ratio of the measured data is 30. The input (*u*_1_) is given as unit-impulse given at *t* = 0, and the model is simulated for *t* = 1 to *t* = 10 seconds. Parameters are estimated using Markov Chain Monte Carlo-based Approximate Bayesian Computation (ABC-MCMC).

From Fig. 8a and Fig 8b, the information gain index *β* and the prediction uncertainty measure *E*(*ζ*) are maximum and minimum, respectively, for the true model (*M*_1_) from which the data is generated. Thus, the proposed method has selected the best model that explains the data in terms of parameter and prediction uncertainty.

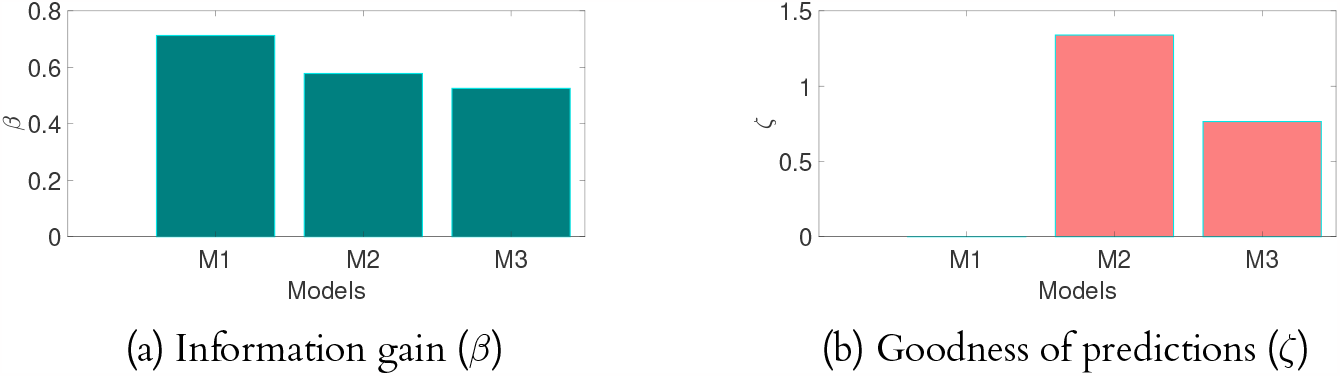

Table 7 contains the parameter estimates and standard errors for all the candidate models. Also, from Figs 9a and 9b, we learn that the parameters *k*_01_ and *k*_21_ are sensitive/stiff parameters and *k*_02_ and *k*_12_ are sloppy/insensitive parameters. The proposed method has chosen the model with low parameter bias and uncertainty on stiff parameters. It can also be seen that the point estimates of the stiff parameter *k*_21_ in *M*_2_ have a high bias. Similarly, the point estimates of *k*_01_ and *k*_21_ in *M*_3_ have high bias.

**Table 7:** Parameter estimates of all the candidate models.

| Models | $k_{01}$ | $k_{02}$ | $k_{12}$ | $k_{21}$ |
| --- | --- | --- | --- | --- |
| $M_1$ | 0.50 ($\pm 0.05$) | 1.59 ($\pm 0.22$) | 2.09 ($\pm 0.22$) | 1.49 ($\pm 0.05$) |
| $M_2$ | - | 1.58 ($\pm 0.18$) | 2.09 ($\pm 0.19$) | 2.65 ($\pm 0.02$) |
| $M_3$ | 0.83 ($\pm 0.051$) | - | 2.10 ($\pm 0.19$) | 2.48 ($\pm 0.07$) |

**Table 8:**
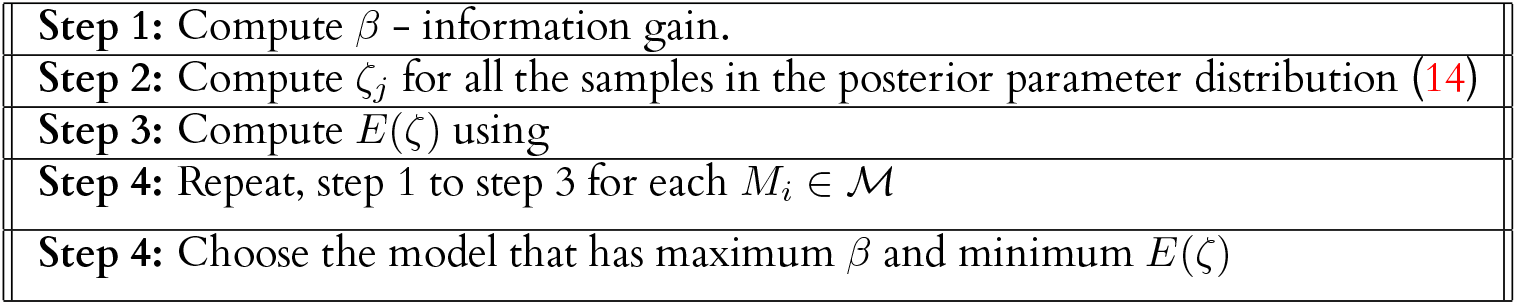
Procedure for model selection.

|  |
| --- |
| **Step 1:** Compute $\beta$ - information gain. |
| **Step 2:** Compute $\zeta_j$ for all the samples in the posterior parameter distribution (14) |
| **Step 3:** Compute $E(\zeta)$ using |
| **Step 4:** Repeat, step 1 to step 3 for each $M_i \in \mathcal{M}$ |
| **Step 4:** Choose the model that has maximum $\beta$ and minimum $E(\zeta)$ |

**Figure 9:**
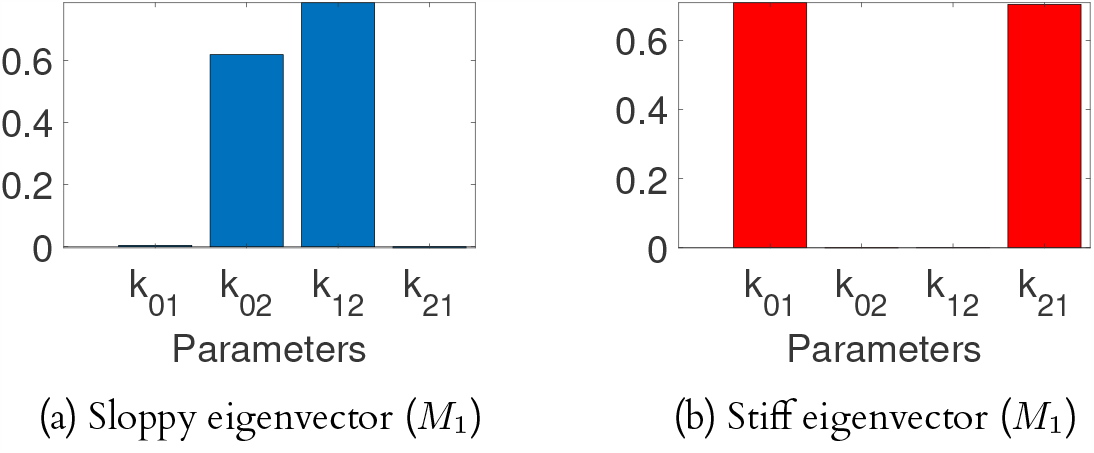
Sloppy and stiff eigenvectors

Fig. 10 contains predictions from sample posterior distributions generated from each candidate model. The predictions from *M*_2_ and *M*_3_ are highly uncertain and have a significant systematic error compared to *M*_1_. The bias in the estimates of the stiff parameters is the prime reason. Hence, the proposed model selection method selects the best model based on the goodness of predictions and parameter estimates. Maximising *β* information gain index for joint distributions minimises the variance and bias of stiff (sensitive) parameters. Consequently, The prediction uncertainty is tightly bounded around the data points.

**Figure 10:**
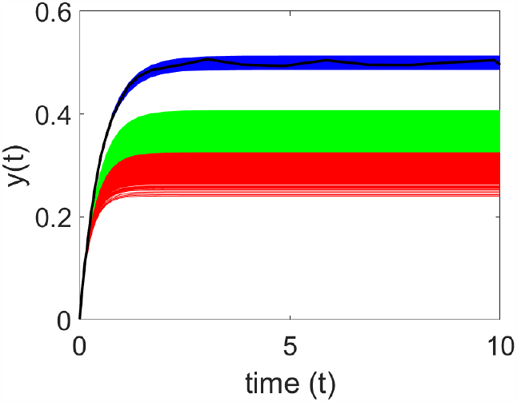
Comparison of prediction uncertainty of candidate models; Blue - predictions from sample posterior of model (*M*_1_). Green - predictions from sample posterior of model (*M*_2_). Red - predictions from sample posterior of model (*M*_3_). Black - data (*z*).

**Figure 11:**
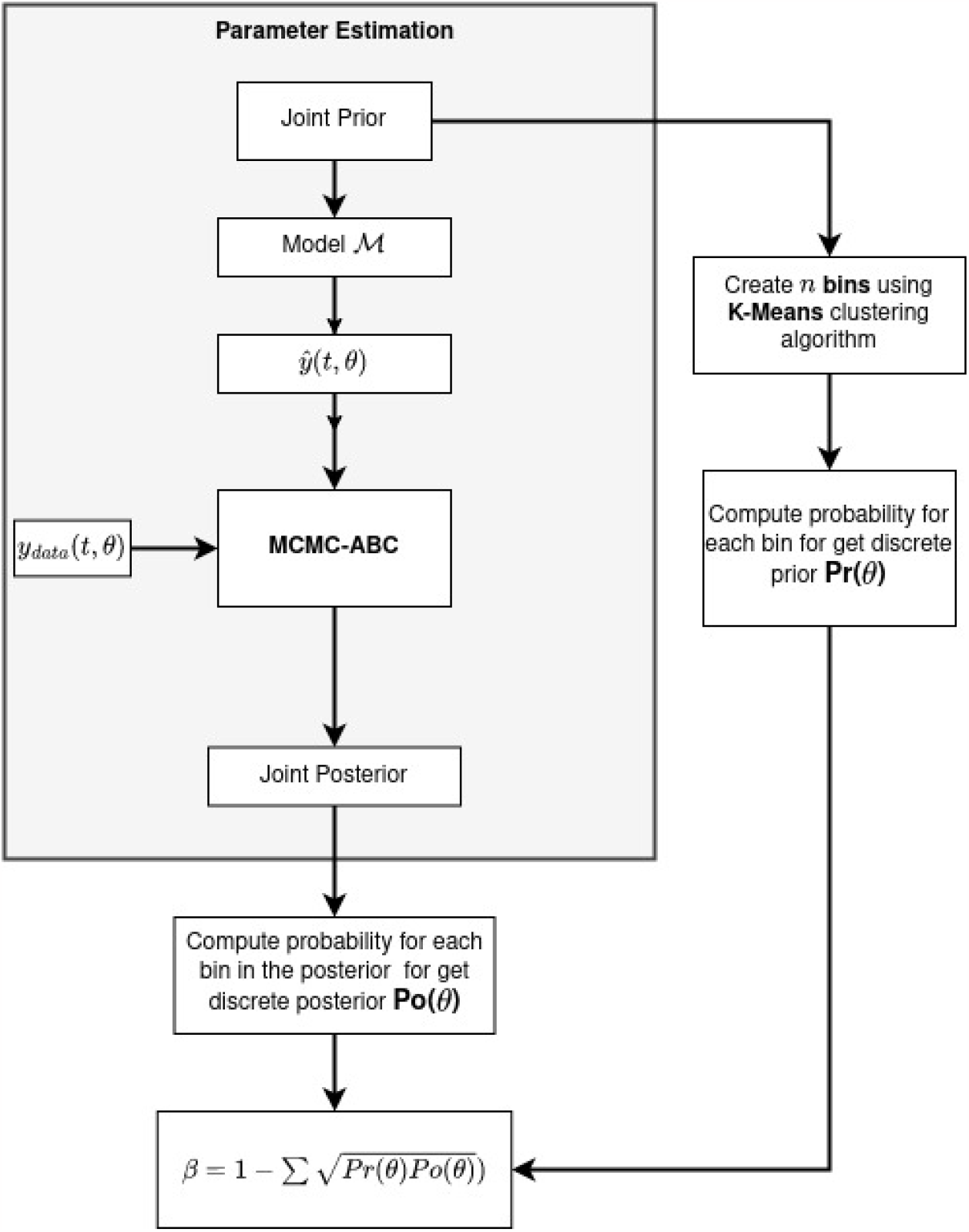
Algorithm for estimating *β* information gain for generalised prior and posterior distributions

## 4 Methodology

In this section, we discuss the procedure for Bayesian Optimal Experiment Design and model selection for generalised priors. Given a model *M*, a prior of the parameters in the form of probability distribution and a set of experimental conditions 𝒳 = *z*_1_, *z*_2_, · · ·, *z*_*n*_, the aim is to select the experiment condition *z*_*i*_ that maximises the *β*-information gain.

### 4.1 Procedure for estimating *β*-information gain

In order to estimate the Bhattacharyya coefficient for discrete distributions, we need to create bins in the parameter space and estimate the PMF. In this work, we employ the *K*-means algorithm to create bins in the parameter space. We choose the number of bins for each parameter as 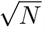, where *n* is the number of samples in the prior parameter distribution. The procedure for estimating the new information gain is given below

#### Procedure

1. Sample *N* points from joint prior distribution *P* (*θ*)
2. Compute the number of bins using

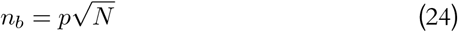

where *n*- dimension of the parameter space.
3. Create *n*_*b*_ number of bins in the prior parameter space using *k*-means clustering algorithm.
4. Estimate the PMF of the prior *P* (*θ*) by computing the probability for each bin.
5. Initialise *θ*_*i*_, *θ*_*i*_ = 1
6. Propose *θ*^*^ from prior distribution *P* (*θ*)
7. Simulate a data set *z*^*^ from the model ℳ(*θ*^*^)
8. If *d*(*zd*_*d*_, *z*^*^) *< ϵ*, go to Step 9, else set *θ*_*i*+1_ = *θ*_*i*_ and go to Step 10.
9. Set *θ*_*i*+1_ = *θ*^*^ with probability

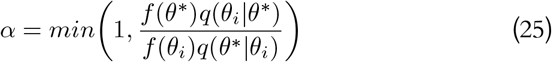

and *θ*_*i*+1_ = *θ*_*i*_, with probability 1 − *α*
10. Accept *θ*^*^ into posterior *P* (*θ*|*zd*_1_), increment *i* = *i* + 1 and go to Step 6
11. Estimate the PMF of the posterior distribution *P* (*θ*|*zd*_*i*_)
12. Estimate the *β*-information gain using Eq. 6

We use Markov Chain Monte Carlo-based Approximate Bayesian Estimation (MCMC-ABC) to estimate the posterior distribution of parameters [35]. The plain ABC rejection algorithm will have a low acceptance rate when the prior distribution of parameters is different from the true posterior distribution.

### 4.2 Procedure for model selection

As a first step, for each model structure *M*_*i*_ ∈ *M*, we estimate the *β* information gain index between the joint prior and posterior of the parameter estimates. From (13), it is evident that maximising the information gain index *β* minimises the uncertainty in the parameter estimates. This gain in information indicates the practical identifiability of parameter estimates for each model *M*_*i*_. Further, we compute the goodness of predictions using (15).

*E*(*ζ*) is estimated using sample mean. The model with minimum *E*(*ζ*) has the least prediction uncertainty. Thus, the model that results in the least prediction and parameter uncertainty is the winner. However, when the model set does not contain the true model that generated the data, a single model resulting in low predictions and parameter uncertainty is not guaranteed; in such scenarios, the user can choose a model with a trade-off between prediction and parameter uncertainty.

## 5 Discussion

The design of optimal experiments is one of the crucial aspects in the system identification of biological systems, primarily due to the difficulty and cost associated with conducting experiments. In this work, we extend our previous work [30] on Bayesian Optimal Experiment Design to the generalised parameter distributions and their application in model selection.

Firstly, using the numerical Bayesian estimation algorithm, the priors and posteriors are not always analytically tractable distributions, except in a few special cases. Hence, we estimate the *β*-information gain using the discrete version of the Bhattacharyya coefficient. The proposed version of the *β*-information gain has a beautiful geometric interpretation; maximising the information gain is equivalent to increasing the angle between the prior and posterior PMFs. Further, we show that for a uniform prior (uninformative) and Kronekar delta (deterministic) posterior with countably infinite sample space, the angle between prior and posterior is *π/*2. Further, we propose a method to estimate the discrete version of *β*-information gain.

We demonstrated the working of our method in two realistic experiment design problems with high relevance in systems biology. Firstly, we used the proposed method to choose the best measurement function (implicitly measurement method) for precise and accurate parameter estimation in Hes 1 transcription network. The *β*-information gain chose the experiment with minimum mean square error, and all the stiff parameters were estimated with minimum bias and variance. Secondly, we employed the proposed method to select the optimal six-point sampling schedule for the HIV1 2-LTR dynamics model. Given the regulations and costs of clinical trials, this problem is paramount in healthcare. Out of four sampling schedules considered, the *β* information gain again chose the six-point sampling schedule, resulting in a minimum mean square error of parameter estimates. Further, the predictions using the parameters from the *β*-optimal resulted in the least prediction uncertainty.

Finally, we proposed a novel method for model selection in systems biology using the *β*-information gain. We demonstrated the working of the method in compartmental models. The proposed method has rightly identified the true model. Thus, we believe that the proposed *β* information gain has vast potential to be applied in experiment design and model selection. The demonstrated examples show that the use of *β*-information gain in the experiment design for systems biology unambiguously chooses experiments that result in minimum parameter and prediction uncertainty. Hence, we believe that generalized *β*-information gain has significant utility in several aspects of system identification, such as OED and model selection.

## Data Availability

The link for the code for estimating *β*-information gain is provided in the supporting information file 2.

## Authors’ Contributions

**Conceptualization**: Karthik Raman, Arun K. Tangirala

**Formal analysis**: Prem Jagadeesan

**Funding acquisition**: Karthik Raman, Arun K. Tangirala

**Investigation**: Prem Jagadeesan

**Methodology**: Prem Jagadeesan, Karthik Raman, Arun K. Tangirala

**Supervision**: Karthik Raman, Arun K. Tangirala

**Validation**: Prem Jagadeesan

**Writing-original draft**: Prem Jagadeesan

**Writing - review and editing**: Prem Jagadeesan, Karthik Raman, Arun K. Tangirala

## 6 Supporting information

### 6.1 HIV 1 2-LTR Dynamics

**Figure 12:**
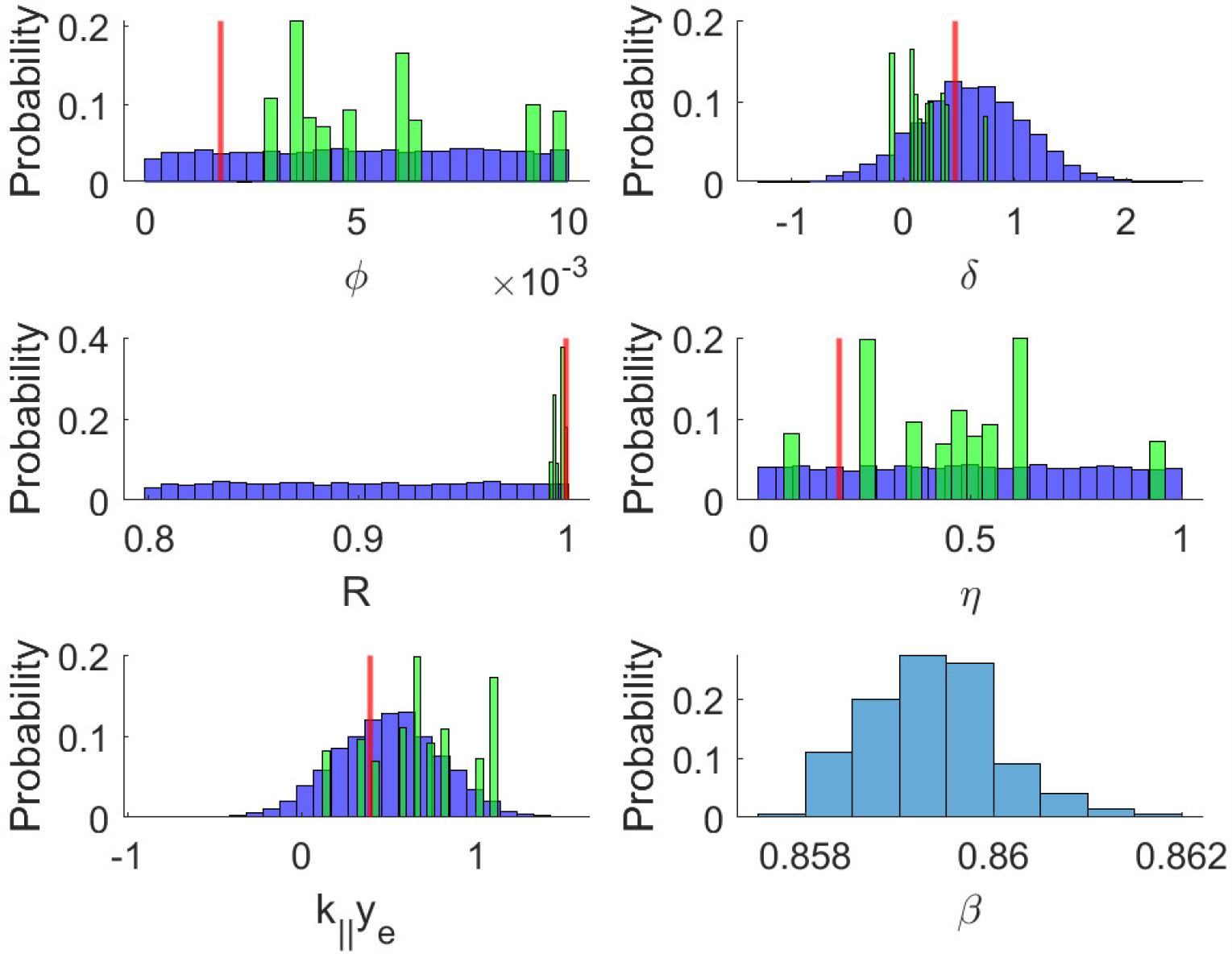
Sample prior and posterior parameter distributions for *A*-optimal sampling schedule. Blue-sample prior distribution, Green-sample posterior distribution, Red-true parameter

**Figure 13:**
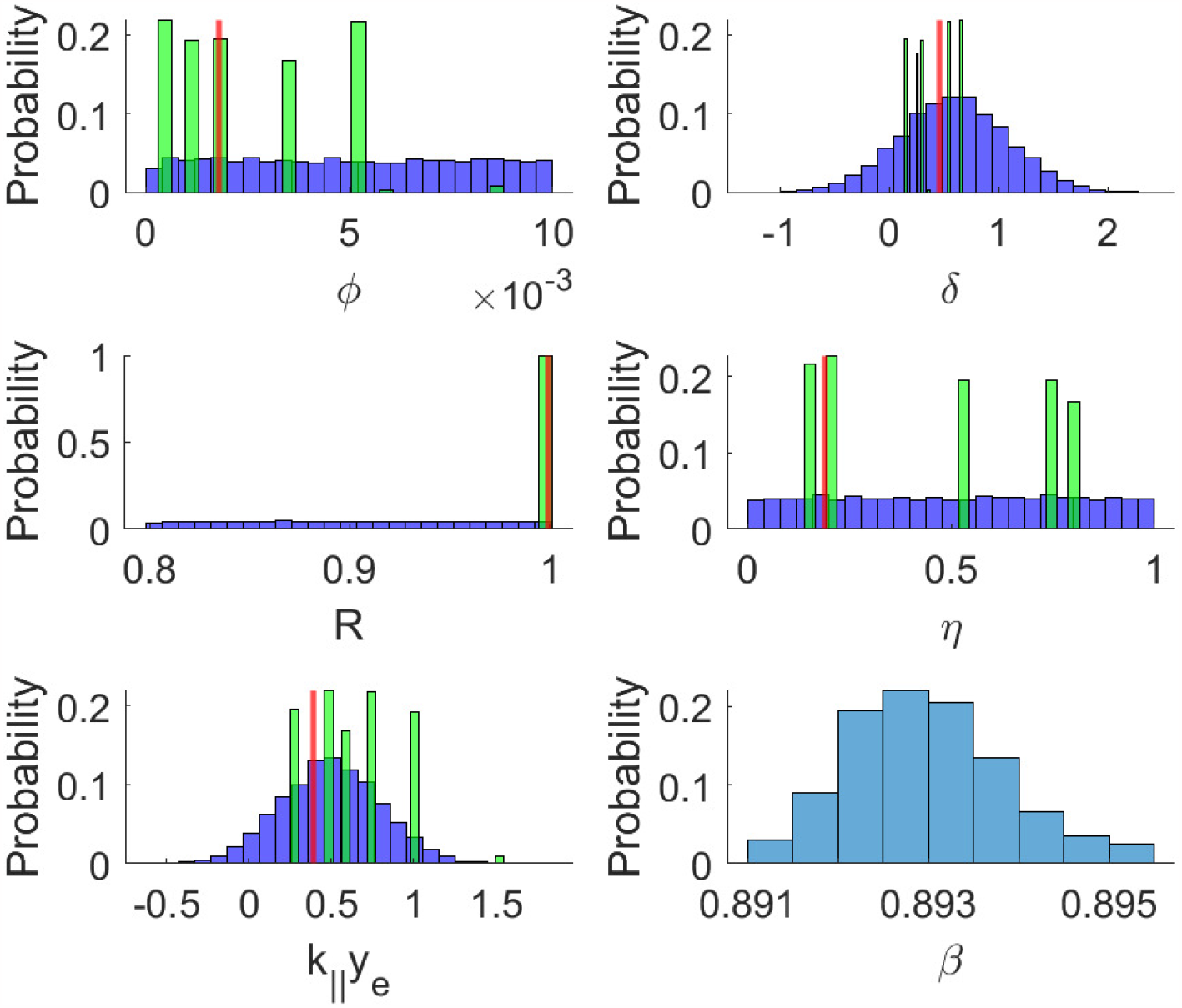
Sample prior and posterior parameter distributions for *D*-optimal sampling schedule. Blue-sample prior distribution, Green-sample posterior distribution, Red-true parameter

**Figure 14:**
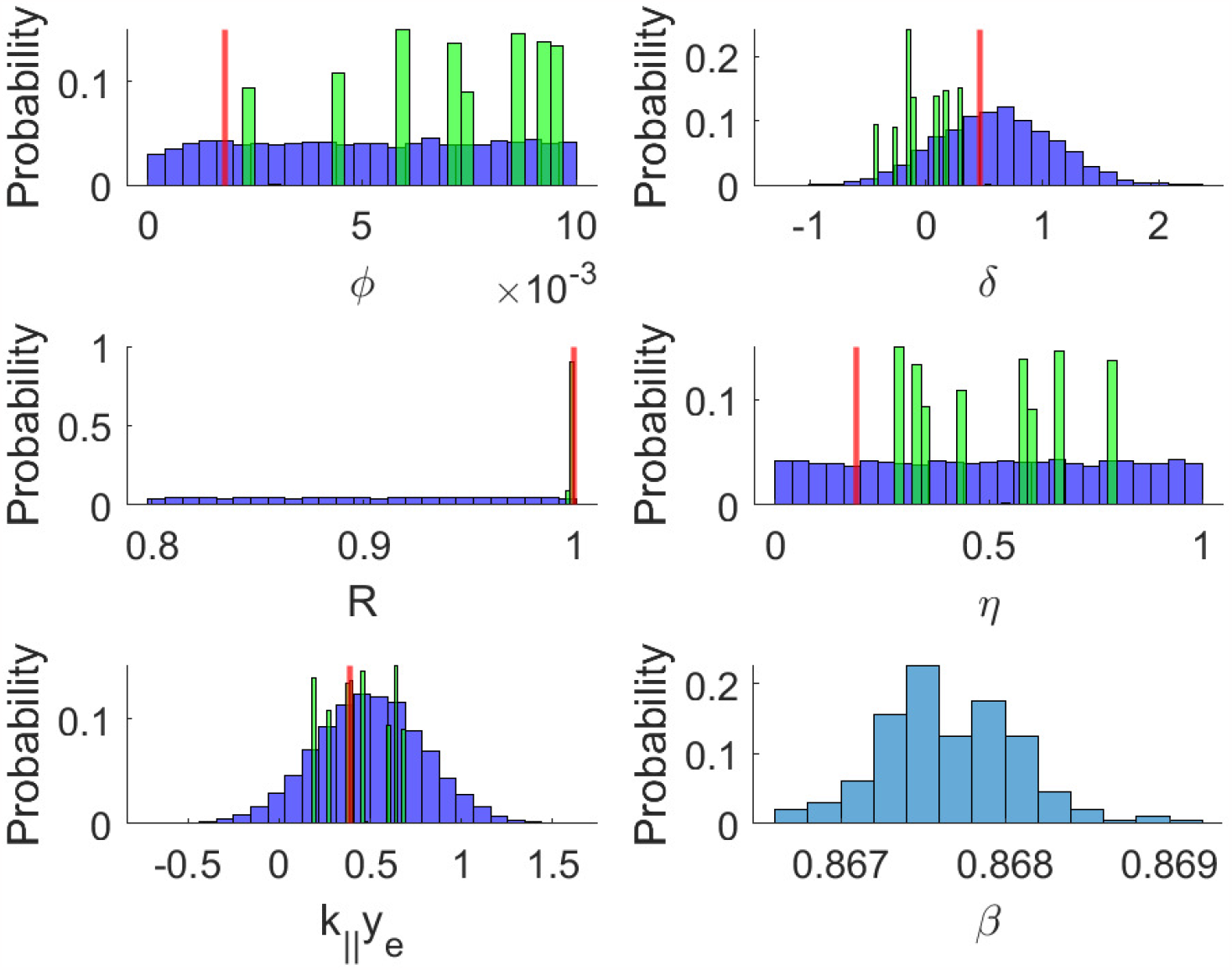
Sample prior and posterior parameter distributions for *E*-optimal sampling schedule. Blue-sample prior distribution, Green-sample posterior distribution, Red-true parameter

**Figure 15:**
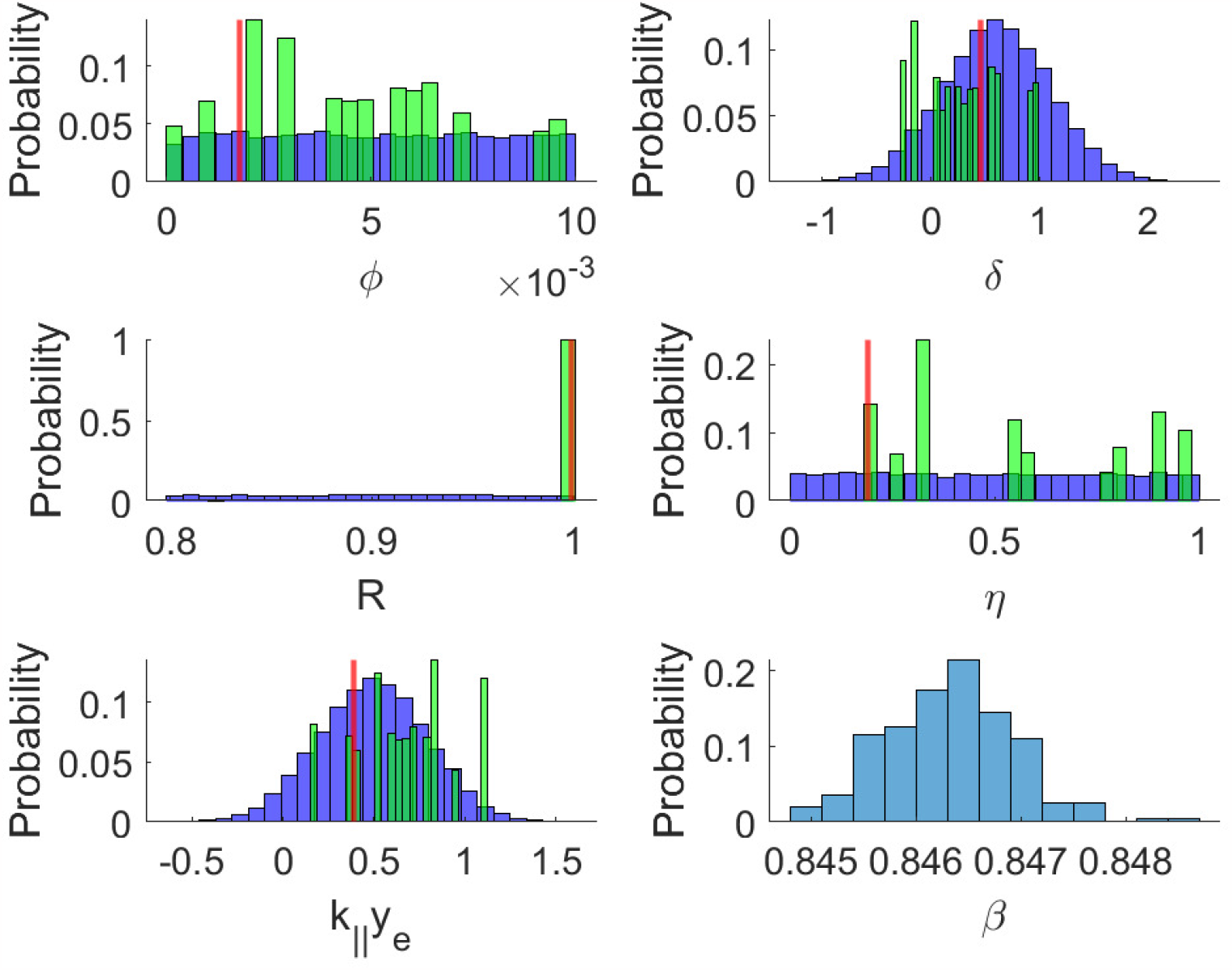
Sample prior and posterior parameter distributions for *ELK*-optimal sampling schedule. Blue-sample prior distribution, Green-sample posterior distribution, Red-true parameter

